# Deciphering the Wisent Demographic and Adaptive Histories from Individual Whole-Genome Sequences

**DOI:** 10.1101/058446

**Authors:** Mathieu Gautier, Katayoun Moazami-Goudarzi, Leveziel Hubert, Hugues Parinello, Cécile Grohs, Stéphanie Rialle, Rafal Kowalczyk, Laurence Flori

**Affiliations:** CBGP, INRA, CIRAD, IRD, Supagro, 34988 Montferrier-sur-Lez, France; IBC, Institut de Biologie Computationnelle, 34095 Montpellier, France; GABI, INRA, AgroParisTech, Université Paris-Saclay, 78350 Jouy-en-Josas, France; UGMA, INRA, Université de Limoges, 87060 Limoges, France; MGX-Montpellier GenomiX, c/o Institut de Génomique Fonctionnelle, Montpellier, France; Mammal Research Institute, Polish Academy of Sciences, 17-230 Białowieża, Poland; INTERTRYP, CIRAD, IRD, 34398 Montpellier, France

## Abstract

As the largest European herbivore, the wisent (*Bison bonasus*) is emblematic of the continent wildlife but has unclear origins. Here, we infer its demographic and adaptive histories from two individual whole genome sequences via a detailed comparative analysis with bovine genomes. We estimate that the wisent and bovine species diverged from 1.7×10^6^ to 850,000 YBP through a speciation process involving an extended period of limited gene flow. Our data further support the occurrence of more recent secondary contacts, posterior to the *Bos taurus* and *Bos indicus* divergence (*ca*. 150,000 YBP), between the wisent and (European) taurine cattle lineages. Although the wisent and bovine population sizes experienced a similar sharp decline since the Last Glacial Maximum, we find that the wisent demography remained more fluctuating during the Pleistocene. This is in agreement with a scenario in which wisents responded to successive glaciations by habitat fragmentation rather than southward and eastward migration as for the bovine ancestors.

We finally detect 423 genes under positive selection between the wisent and bovine lineages, which shed a new light on the genome response to different living conditions (temperature, available food resource and pathogen exposure) and on the key gene functions altered by the domestication process.

## Introduction

The European bison, *Bison bonasus* (BBO), also known as the wisent, is an emblematic species of the European wildlife. The oldest fossil remains were found in Eastern Europe and trace back to the Late Pleistocene, approximately 12,000 Years Before Present (YBP) (Benecke 2005). The wisent is closely related to the North American bison (*Bison bison*), both species assumed to be derived from the extinct long-horned steppe bison *(Bison priscus)* which was widespread across the Northern hemisphere in the mid and upper Pleistocene (i.e., 781,0 to 11,700 YBP). However, several morphological and behavioral traits distinguish the wisent from its American relative. In particular, adults have generally a smaller body length (Krasińska and Krasiński 2002), display a less hairy body and tend to be mixed feeders that combine browsing and grazing (Kowalczyk, et al. 2011; Merceron, et al. 2014) while American bisons are essentially grazer. These various characteristics might be viewed as adaptation to living constraints associated to forest habitat that has been considered as a refuge (Kerley, et al. 2012; Bocherens, et al. 2015). During the mid and late Holocene, the wisent was distributed in central and eastern Europe with a longest survival in north-eastern Europe (Benecke 2005). Nevertheless, the progressive replacement of the open steppe by forest cover in early Holocene and the human population growth associated with the spread of farming during the middle Holocene lead to a reduction of its habitat (Kuemmerle, et al. 2012; Bocherens, et al. 2015). Hence, by the Middle Age, the wisent was only present in a few natural forests of North-Eastern Europe and finally faced extinction in the wild at the beginning of the 20^th^ century (Pucek 2004). A species restoration program relying on a few tens of specimens kept in European zoos and private breeding centers was early initiated in the 1920’s (Pucek 2004; Tokarska, et al. 2011). Thanks to these efforts, the wisent world population now consists of over 5,000 animals (including about 3,500 free-living individuals) that are mostly maintained in forests of eastern Europe (Kerley, et al. 2012; Raczyński 2015). The wisent is among the few large mammalian terrestrial species that survived the massive megafauna extinction of the last glacial/interglacial transition period, ca. 25,000-10,000 YBP (Lister and A.J. 2008; Lorenzen, et al. 2011). Characterizing the specific conditions of its survival may thus help understanding how organisms evolve in response to environmental change. In this study, we explore the wisent demographic and adaptive histories using individual whole-genome sequences from two males of the wisent lowland line (Tokarska, et al. 2011) living in the Bialowieza Forest. To that end, we perform a detailed comparative analysis of these newly generated wisent genome sequences with individual genome sequences representative of different European cattle breeds. Domestic cattle, the wild ancestor of which, the aurochs (*Bos primigenius*) went extinct in the 17^th^ century, is indeed among the closest wisent relatives (Verkaar, et al. 2004) and benefits from abundant genomic resources (Liu, et al. 2009). Our genome-wide comparison provides an accurate characterization of the divergence between the bovine and wisent lineages and an estimation of their population size histories during Pleistocene and Holocene. In addition, the identification and functional annotation of genes under positive selection between both lineages shed light into the biological functions and the key adaptive traits that were affected by environmental constraints.

## Results and discussion

### Comparison of wisent and cattle genomes reveals low amount of nucleotide divergence

High-throughput sequencing data from two wisent bulls, namely BBO_3569 and BBO_3574, were each aligned onto the UMD3.1 cattle (BTA) genome assembly (Liu, et al. 2009). The read mapping statistics showed overall good performances (Table S1) with noticeably 97.4% (respectively 96.6%) of the reads being properly paired for the BBO_3569 (respectively BBO_3574). Conversely, following the same read mapping procedure but considering as a reference the *OviAr3* genome assembly (Jiang, et al. 2014) from the more distantly related domestic sheep (OAR) strongly altered the statistics (Table S1). Altogether, these mapping results provided support to consider the cattle genome as closely related enough to provide a good reference to assemble our wisent reads. For autosomes, an approximately 10X-coverage was achieved for both individuals (9.81X and 11.6X for BBO_3569 and BBO_3574 respectively) with more than 95% of the (bovine) reference sequence covered by at least one read (Table 1). As expected for males, the bovine chromosome X was only slightly more than half covered (ca. 56.8% and 55.8% of the autosome coverage for BBO_3569 and BBO_3574 respectively) due to the inclusion of pseudo-autosomal regions (PAR) in the estimation. Conversely, the (bovine) mitochondrial genome was found highly covered (>300X) for both individuals.

**Table 1:** Read mapping statistics from the alignment of each BBO_3569 and BBO_3574 European bison genome sequences onto the UMD 3.1 cattle genome assembly (Liu, et al. 2009).

| Chromosome Type (BTA) | Size (in bp) | Coverage (% of sequence covered)* |  | Average nucleotide divergence in % (number of sites compared)** |  |
| --- | --- | --- | --- | --- | --- |
|  |  | BBO_3569 | BBO_3574 | BBO_3569 | BBO_3574 |
| Autosomes | 2,512,082,506 | 9.81 (95.5) | 11.6 (95.7) | 0.870<br>(2.27x10<sup>9</sup>) | 0.882<br>(2.29x10<sup>9</sup>) |
| X-chromosome (including PAR) | 148,823,899 | 5.57 (90.9) | 6.48 (91.2) | 0.636<br>(1.11x10<sup>8</sup>) | 0.642<br>(1.17x10<sup>8</sup>) |
| Mitochondria | 16,338 | 397 (97.7) | 302 (97.1) | 3.58<br>(1.18x10<sup>4</sup>) | 3.41<br>(1.20x10<sup>4</sup>) |

\* For each individual, the average read coverages (after alignment) over autosomes, the X-chromosome and the mitochondria are given together with the overall percentages of sites from the corresponding reference sequences covered with at least one read.
\*\* For each type of chromosomes, the average nucleotide divergence between the cattle reference genome and each European bison individual consensus sequence is given (see Material and Methods).

Interestingly, the estimated autosomal nucleotide divergence between the bovine genome and each wisent individual genome was found equal to 0.870% for BBO_3569 and 0.882% for BBO_3574 (Table 1) supporting a low amount of genome divergence between the two species. The estimated nucleotide divergence was about 1.37 times smaller for the chromosome X than for autosomes, as already observed in the Chimpanzee vs. Human genome sequence comparison (Mikkelsen, et al. 2005). This is likely related to a higher mutation rate in the male compared to the female germ line in mammalian species (Li, et al. 2002; Mikkelsen, et al. 2005). Also, the nucleotide divergence increased in distal chromosomal regions in both autosomes and X-chromosome comparisons while read coverage remained mostly uniform (Figure S1). Such a regional pattern, also reported in the aforementioned Chimpanzee vs. Human genome sequence comparison (Mikkelsen, et al. 2005), might similarly result from the physical properties of the corresponding distal regions (i.e. high local recombination rate, high gene density and higher GC content). Indirectly, observing such a pattern for wisent sequences mapped onto the cattle genome assembly supports a low amount of chromosome rearrangements between the two species. Finally, the estimated nucleotide divergence for the mitochondrial genomes were about 4 times higher than for autosomes. This suggests an absence of recent maternal bovine introgression at least within the lineages of the two wisents we considered.

### The wisent demography has been more fluctuating than the bovine one during the whole Pleistocene

We further sought to characterize the genetic variability of the wisent population that has long been considered as a threatened species and is still classified as “vulnerable” by the International Union for Conservation of Nature (IUCN) red list (http://www.iucnredlist.org/). The estimated average (autosomal) heterozygosities, as approximated by the population mutation rate (*θ* = 4*N_e_μ_g_* where *μ_g_* represents the mutation rate per site and per generation and *N_e_* the effective population size) measured on whole individual genomes, were consistent when considering either the BBO_3569 (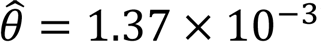) or the BBO_3574 (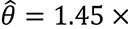 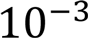) individual. Interestingly, both estimates were similar to the ones obtained on four individual cattle genomes representative of the Angus (AAN_0037), Jersey (JER_0009), Holstein (HOL_0101) and Simmental (SIM_0043) European cattle breeds which ranged from 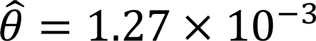 to 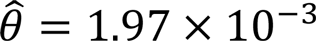 (Figure S2 and Table S2). The small heterozygosity observed for wisent might be explained by the sharp decline of its effective population size during the last 20,000 years (see below), but also by its more recent history since wisent experienced a strong bottleneck at the end of the First World War (Wójcik, et al. 2009; Tokarska, et al. 2011). However, we noted that our wisent estimates remained within the range observed for the four different European cattle breeds. This suggests that the constitution of the Białowieża wisent herd from a small number of presumably poorly related founders and subsequent management (Pucek 2004) allowed to recover a reasonable amount of variability for the current population.

More generally, the patterns of variability observed at a local individual genome scale are informative about the demographic history of the population (Li and Durbin 2011). From our two individual genomes, we estimated the wisent past *N_e_* using the Pairwise Sequentially Markovian Coalescent model (PSMC) introduced by Li and Durbin (2011), and compare it to that estimated for the bovine species based on each of the four cattle individual genomes aforementioned. Estimates of the scaled effective population sizes (in units of 4*N_e_μ_g_*.) backward in time measured in scaled units of 2*μ_y_T* (where *T* is in YBP, and *μ_y_* represents the mutation rate per site and per year) are plotted in Figure S3 for each of the six individual genome analyses (see Figure S4 for bootstrap confidence intervals of each history). Overall, a similar trend was observed within each species, irrespective of the individual considered (although the AAN_0037 profile was more dissimilar than the three other cattle ones probably as a result of an overall lower coverage for this individual). By contrast, marked differences were found when comparing the inferred wisent and bovine population size histories. In particular, from backward time (in scaled units) *t=10^−3^* to *t=5.10^−5;^* (Figure S3), wisent displayed more pronounced oscillations and from twice to three times higher effective population sizes. Because the PSMC model relies on local density of heterozygosites, using the bovine genome assembly as a reference to derive the analyzed individual wisent genomes (see Material and Methods) could have lead to some inferential biases (Nevado, et al. 2014). Nevertheless, the inferred wisent population size histories remained similar when considering a BBO consensus genome as a reference to derive the individual genome sequences (Figure S5). Similarly, given the close relatedness of the bovine and bison species, chromosomal rearrangements might remain negligible and should not have substantial effect on the overall inferred histories.

To facilitate biological interpretation of the observed demographic signals, we transformed the population size history profiles assuming *μ_y_* = 2.10 × 10^−9^ (Liu, et al. 2006) to translate the time scale in YBP. Further, to translate the effective population sizes (*N_e_*) in real units, we assumed a generation time of *g*=6 years (Keightley and Eyre-Walker 2000; Gautier, et al. 2007), leading to *μ_g_* = *g* × *μ_y_* = 1.26 × 10^−8^. The resulting backward in time estimates of *N_e_* are plotted in Figure 1. For both the wisent and bovine species, a sharp decline in *N_e_* started at the end of the last glacial period, predating cattle domestication approximately 10,000 YBP as already suggested for the latter species by MacLeod *et al.* (2013). This most presumably originates from a growing impact of human activities via hunting intensification and land anthropization especially after the development of agriculture (Soares, et al. 2010). Interestingly, the estimated wisent population sizes clearly oscillated according to the succession of glacial Pleistocene periods to *ca*. 500,000 YBP (viewed backward in time) with interglacial periods coinciding with lower population sizes. Conversely, from the Last Glacial Maximum (LGM) approximately 20,000 YBP (Clark, et al. 2009) to 150,000 YBP, the estimated bovine population sizes only slightly decreased, remaining always between 10,000 to 20,000 individuals (more than two to three times lower than wisents, as mentioned above). Hence, even if the fluctuating pattern observed for wisent might correspond to actual changes in the population sizes, the comparison with the bovine species might lead to an alternative scenario of population fragmentation. In such a scenario, the alternation of population isolation events due to fragmentation of the habitats during glacial periods leads to an increase of the overall estimated Ne while a more continuously interbreeding between populations during interglacial periods leads to a decrease in the overall estimated Ne. Indeed, as recently evidenced by Mazet *et al.* (2015) when considering structured populations, changes in the gene flow patterns result in changes in the estimated *N_e_* (even if the actual *N_e_* remained constant) as PSMC assumes a panmictic model. As a result, one might expect an increase (respectively a decrease) in the estimated *N_e_* when gene flow and thus the amount of connectivity between (sub)populations decreases (respectively increases). It is not possible from the PSMC analyses to distinguish between a scenario involving habitat fragmentation and a scenario of actual population size changes. However, it is tempting to speculate from the divergent patterns in the inferred population size across the wisent and bovine species that the ancient aurochs distribution range, possibly restrained to southern regions with more stable habitats, has only been partially overlapping the wisent one (hence the lower *N_e_*). In agreement with the “refugium theory” (Hofreiter and Stewart 2009), both species might have followed different strategies to survive during Pleistocene glaciations, the wisent remaining in small disconnected refugia while the aurochs migrated southward and possibly eastward (Mona, et al. 2010). Conversely, during interglacial periods, wisent may have retrieved a larger distribution area with a reconnection of isolated populations while the aurochs may have recolonized northern and central areas. Accordingly, the divergence of the wisent and the American bison which took place *ca*. 230,000 YBP (Hassanin, et al. 2013), might result from an absence of reconnection between populations separated in Eurasian and North-American refugia. Note also that the most recent common ancestor of all the Beringian bison lived from 164,000 to 111,000 YBP (Shapiro, et al. 2004), a period that lies within the second peak observed in our estimated wisent population size history (Figure 1).

**Figure 1.**
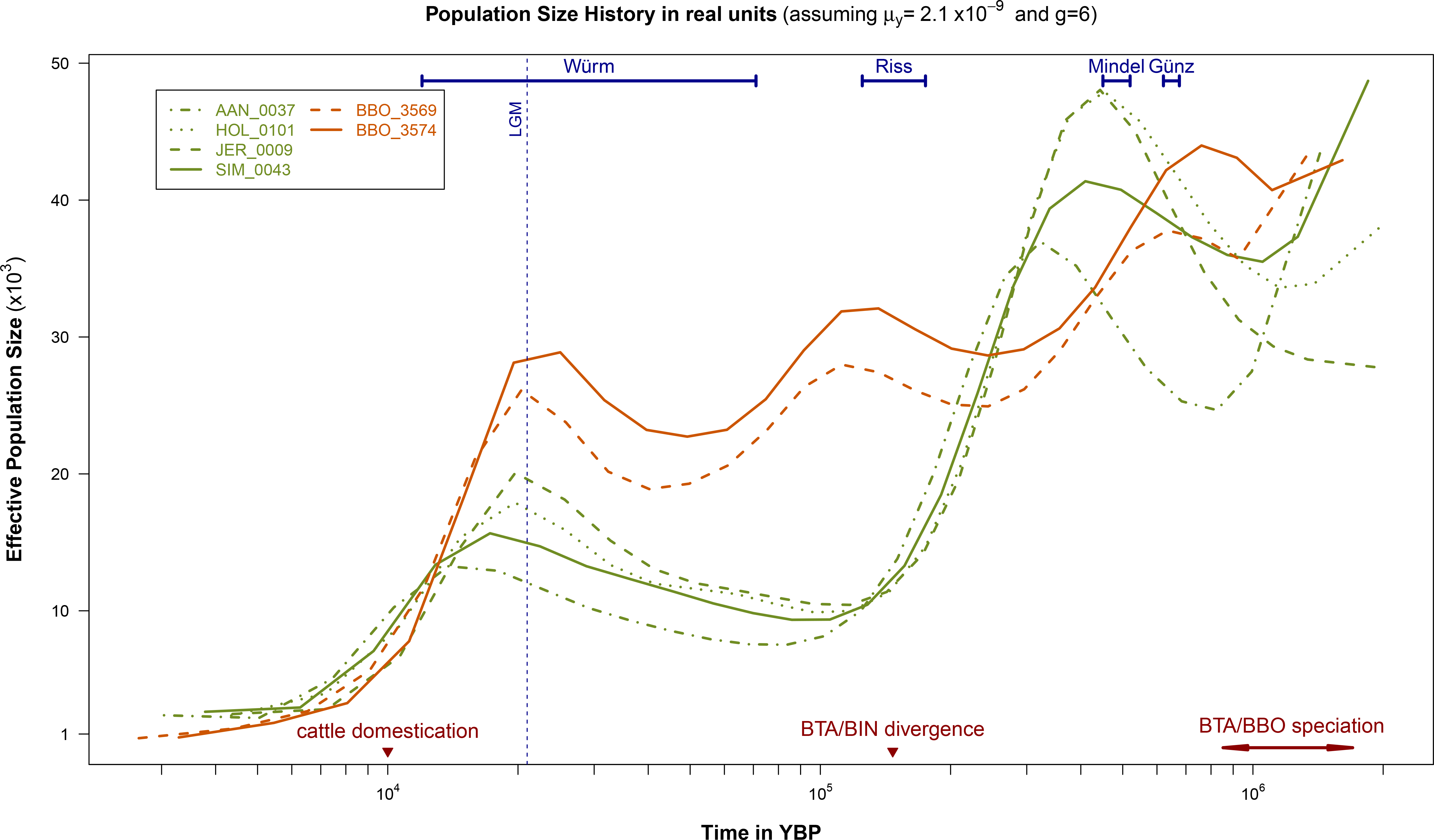
Population size histories inferred from the two wisent and the four bovine genomes under the PSMC model. Backward in time (in YBP) estimates of the effective population sizes derived from the *psmc* analyses of the BBO_3569 and BBO_3574 individual wisent genomes and the AAN_0037, HOL_0101, JER_0009 and SIM_0043 bull bovine genomes and assuming ***μ_y_* = 2.10 × 10^−9^** (Liu, et al. 2006) and *g*=6 (Keightley and Eyre-Walker 2000; Gautier, et al. 2007). At the bottom of the figure, the timing of cattle domestication and the zebu (BIN) and taurine (BTA) divergence (Ho, et al. 2008) are indicated by a dark red triangle while the timing interval of the bovine and wisent divergence (see the main text) is indicated by dark red arrows. Similarly, European ice ages (Würm, Riss, Mindel and Günz glacial stages) according to the Penck and Bruckner work (Penck and Brückner 1901-1909) cited in (Elias 2013) are indicated by blue intervals at the top of the figure. Finally, the Last Glacial Maximum (LGM), ca. 20,0 YBP (Clark, et al. 2009), is indicated by a vertical blue dotted line.

Finally, in agreement with MacLeod et al. (2013) that used a different inference model (and a different time calibration procedure), we also observed a steep increase (viewed backward in time) starting ca. 150,000 YBP in the inferred bovine population sizes that might actually be explained by the divergence of the taurine (BTA) and zebu (BIN) lineages (see e.g., Figure S5 in Li and Durbin, 2011). Indeed, this divergence event is dated between 117,000 and 275,000 YBP based on diversity of cattle mitochondrial sequences (Bradley, et al. 1996) and at 147,0 YBP (with a 95% Highest Posterior Density interval ranging from 84,000 to 219,000 YBP) using modern and radiocarbon-dated ancient mitochondrial DNA (Ho, et al. 2008). A similar increase starting about 1 MYA could also be observed for the ancestral wisent population size which coincides with the timing of BTA/BBO speciation (see below).

### The wisent and bovine species diverged in the early Pleistocene and experienced more recent secondary contacts

To characterize the divergence between the BBO and BTA lineages, we further relied on the coalescent Hidden Markov Models (CoalHMM) framework (Hobolth, et al. 2007). Two models were considered corresponding to *i)* an isolation (I) model which assumes that the ancestral population split into two populations at time *τ_D_* into the past (Mailund, et al. 2011); and *ii)* an isolation-with-migration (IM) model which assumes that the ancestral population started to split into two populations at time *τ_2_*=*τ_1_*+*τ_M_* and that these two populations exchanged gene with a migration rate *M* during a period *τ_M_* until a later time *τ_1_* when gene flow stopped (Mailund, et al. 2012). Under the I-model and assuming as above *μ_y_* = 2.10 × 10^−9^, the median estimate of *τ_D_* (over 10 Mbp segments) was found equal to 1.20 × 10^6^ YBP (ranging from 1.02 × 10^6^ to 1.36 × 10^6^) and 1.23× 10^6^ YBP (ranging from 1.01 × 10^6^ to 1.51× 10^6^) when considering the alignments onto the UMD 3.1 bovine genome assembly of the BBO_3569 and BBO_3574 whole genome sequences respectively (Figure 2A). As expected, the *τ_D_* estimates were always intermediate between the estimated start (*τ_2_*) and end (*τ_1_*) times of the divergence period obtained under the IM-model (Figure 2B). Hence, when considering the BBO_3569 (respectively BBO_3574) sequence the median estimates of *τ_1_* and *τ_2_* allowed to define a period ranging from 0.846 × 10^6^ YBP to 1.70 × 10^6^ YBP (respectively from 0.864 × 10^6^ YBP to 1.72× 10^6^ YBP) for the divergence of the BBO and BTA lineages. During the corresponding divergence periods, the estimated number of migrants (per generation) was found limited with median values of 0.56 and 0.61 for the BBO_3569 and BBO_3574 analyses respectively (Figure 2C). It should also be noticed that the ancestral population sizes estimated under both the I-model (median values of 62,120 and 65,740 for the BBO_3569 and BBO_3574 analyses respectively) and the IM-model (median values of 50,750 and 53,430 for the BBO_3569 and BBO_3574 analyses respectively) were in agreement with those obtained above with the PSMC for the corresponding time period (Figure 1). Finally, as shown in Figure 2D, the comparison of the I-and IM-models of speciation provided a clear support in favor of the IM-model. From these CoalHMM analyses, we thus conclude that the BBO and BTA lineages diverged in the early Pleistocene (from 1.7×10^6^ to 850,000 YBP), in agreement with previous studies that relied on alternative time calibrations (Bradley, et al. 1996; Troy, et al. 2001). Moreover, the divergence between these two lineages involved an extended period of limited gene flow similar to the speciation process previously reported in great apes (Mailund, et al. 2012).

**Figure 2.**
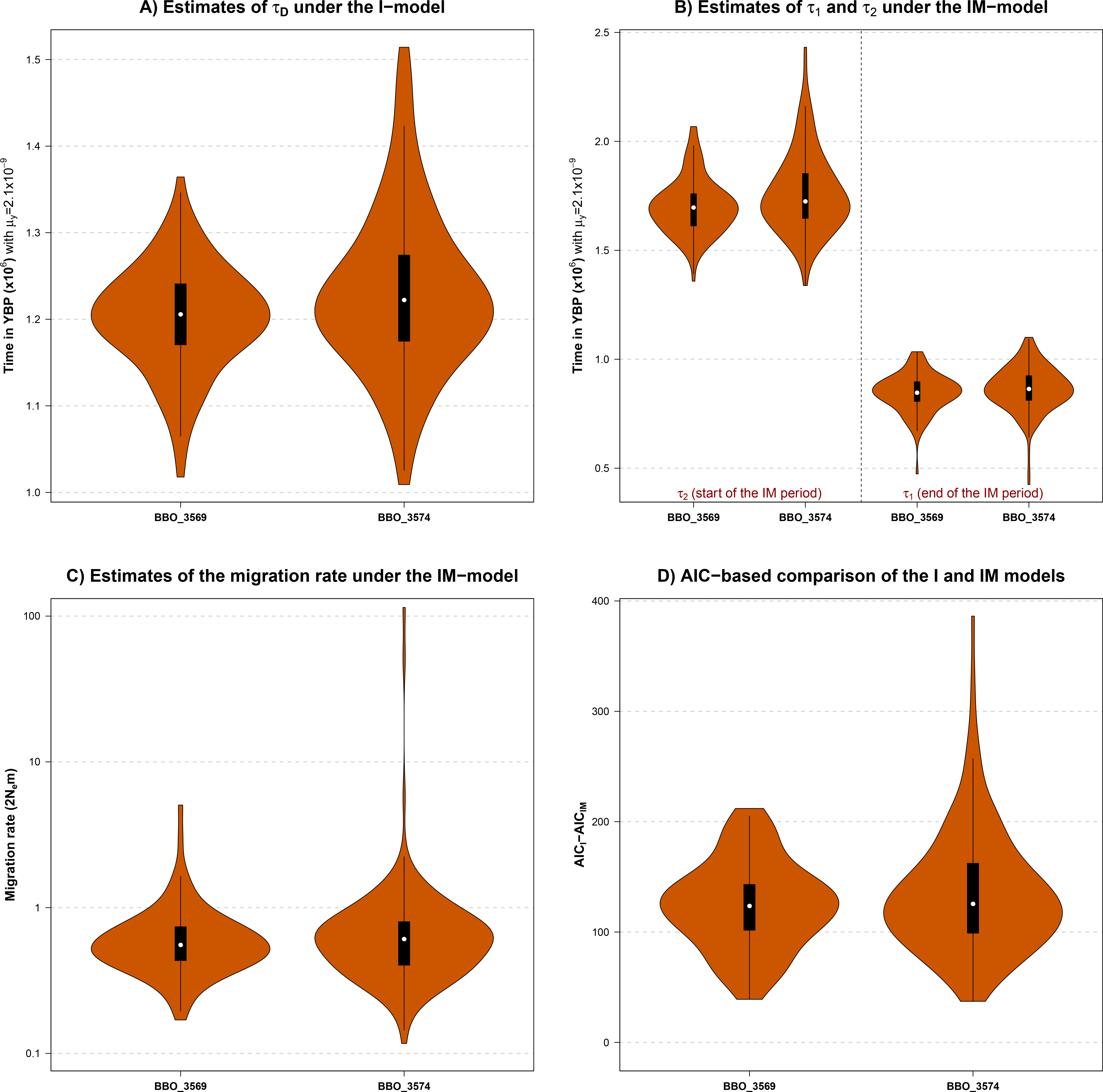
Characterization of the divergence between the BBO and BTA lineages under the isolation and the isolation-with-migration models. A. Estimation of the time *τ_D_* of divergence between the BBO and BTA lineages under the Isolation model (that assumes a clean split). The two violin plots show the distribution of *τ_D_* estimates obtained for each 10 Mbp non-overlapping segments from the alignments of the BBO_3569 (n=123 segments) and BBO_3574 (n=177 segments) whole genome sequence onto the UMD 3.1 bovine genome assembly. Time scale was translated into YBP assuming *μ_y_* = 2.10 × 10^−9^ (Liu, et al. 2006). B. Estimation of starting (*τ_2_*) and ending (*τ_1_*) time estimates of the divergence between the BBO and BTA lineages under the Isolation-with-Migration model (that assumes the two ancestral lineages exchange migrants between *τ_2_* and *τ_1_*). The violin plots show the distribution of *τ_2_* and *τ_1_* time estimates obtained for each 10 Mbp non-overlapping segments from the alignments of the BBO_3569 (n=123 segments) and BBO_3574 (n=177 segments) whole genome sequence onto the UMD 3.1 bovine genome assembly. Time scale was translated into YBP assuming *μ_y_* = 2.10 × 10^−9^ (Liu, et al. 2006). C. Estimation of the average number of migrant per generation (2*N_e_m*) during the divergence of the BBO and BTA lineages under the Isolation-with-Migration model. The two violin plots show the distribution of the migration rate estimates obtained for each 10 Mbp nonoverlapping segments from the alignments of the BBO_3569 (n=123 segments) and BBO_3574 (n=177 segments) whole genome sequence onto the UMD 3.1 bovine genome assembly. D. Model comparisons between the isolation and the Isolation-with-Migration models. The two violin plots show the distribution of the difference between the Akaike Information Criterion for the isolation (*AICI*) and for the isolation-with-migration (*AIC_IM_*) obtained for each 10 Mbp non-overlapping segments from the alignments of the BBO_3569 (n=123 segments) and BBO_3574 (n=177 segments) whole genome sequence onto the UMD 3.1 bovine genome assembly. Because the model with the smallest *AIC* should be preferred, the distributions provide strong support in favor of the IM-model (*AIC_IM_* is always lower than *AICI*).

However, both the I– and IM models were not designed to capture signals from more recent gene flow events associated to secondary contacts between the bovine and wisent species. To that end, we compared the relative abundance of ABBA and BABA site patterns (Green, et al. 2010; Durand, et al. 2011) defined across the following four taxons: *i)* BBO, *ii)* BTA, *iii) Bos indicus* (BIN), and *iv*) OAR. Assuming a (((BTA;BIN);BBO);OAR) phylogeny (e.g., (Buntjer, et al. 2002; Ho, et al. 2008; Jiang, et al. 2014)), ABBA (respectively BABA) sites are those at which the derived allele (“B”) is shared between the nonsister BIN and BBO (respectively BTA and BBO) lineages whereas BTA (respectively BIN) carries the ancestral allele (“A”), as defined by the OAR outgroup. In the absence of (recent) gene flow, both patterns may only result from incomplete lineage sorting and should be equally abundant in the genome while gene flow events between the BTA and BBO lineages would lead to a significant excess of BABA over ABBA sites (Durand, et al. 2011). Whether considering the BBO_3569 or the BBO_3574 sequencing data to define the BBO reference (see Material and Methods), we found a slight but significant excess of “BABA” (n=49,337 and n=58,032 for BBO_3569 and BBO_3574 respectively) over “ABBA” (n=48,004 and n=56,357 for BBO_3569 and BBO_3574 respectively). Accordingly, the *D-statistic* defined as the normalized difference in the counts of ABBA and BABA sites (Green, et al. 2010; Durand, et al. 2011) was found significantly negative for both the BBO_3569 and BBO_3574 derived consensus sequences with values equal to −1.25% (s.d.=0.032%; Z=39.0) and −1.37% (s.d.=0.031%; Z=44.4) respectively. The corresponding proportion of BBO ancestry into (European) cattle was found equal to 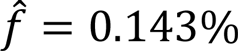 (s.d.=0.004%) which is actually a conservative minimum (Green, et al. 2010). Assuming a constant population size, the bias is equal to *t_a_/t_s_*=0.28/0.85=0.33 in the worst scenario since *t_a_* represents the timing of the BBO and BTA admixture (<275,000 YBP, the lower bound of the estimated divergence between the taurine and zebu lineages, see above) and *t_s_* the timing of the bovine/wisent speciation (>850,000 YBP, the upper bound found above under the IM-model). Even if the resulting corrected proportion 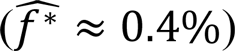 remains very small, the observed (significant) footprints of BBO ancestry into the genomes of European cattle supports the occurrence of secondary contacts between these two lineages that are posterior to the *Bos taurus* and *Bos indicus* divergence *(ca.* 150,000 YBP, see above). Analyzing non-European (e.g., African) taurine genomes might allow to refine the timing of such secondary contacts by assessing whether or not they still occurred after (taurine) cattle domestication (*ca.*, 10,000 YBP). To that respect also, sequencing individuals (or fossil remains) from extant (*Bison bison*, most particularly) or extinct (e.g., *aurochs* and *Bison priscus*) sister species would also be very informative. Conversely, it should be noticed that genomic data from *Bison* species would also allow to quantify the proportion of bovine ancestry (if any) in the BBO genome.

### Identification and functional annotation of genes under positive selection

To identify genes under positive selection between the wisent and bovine lineages, we computed the *Ka/Ks* ratio of nonsynonymous (*Ka*) and synonymous (*Ks*) substitution rates (Kimura 1983) for 17,073 protein sequence alignments between the BBO and BTA genomes. As expected from the effect of purifying selection (Hurst 2002), the average *Ka/Ks* was found equal to 0.273 (ranging from 0.00 to 17.2) with a distribution highly shifted towards 0 (Figure S6). Transcripts with *Ka/Ks>1* were further considered under positive selection leading to the annotation of 425 genes ready to functional and gene network analyses (see Material and Methods, Table S3). Overall, the most significant functions underlying these genes are related to i) nervous system; ii) immune and inflammatory responses; iii) embryonic and organ development; iv) cellular morphology and organization and; v) skeletal and muscular disorders (Table S4). Figure 3 illustrates the connection of the selected genes that were annotated with their key underlying functions and Table 2 gives a more detailed list of functions and sub-functions. Interestingly, many genes are related to several functions (Table S5), suggesting a pleiotropic role and more strikingly, 85% of the genes participated to a global gene network which is defined by six significant networks connected by up to 11 common molecules (Table S6 and Figure S7). Note that very similar results are obtained when restricting the functional analysis to the 359 selected genes that are one to one orthologs between cattle and sheep (see Table 2 and Table S4). Thanks to the method we used to define BBO/BTA gene sequence alignments, this suggests that our annotation remains somewhat robust to extra variation that could have been identified when mapping BBO reads in genic regions belonging to large gene families or due to Copy Number Variants in the sequenced BBO individuals.

**Figure 3.**
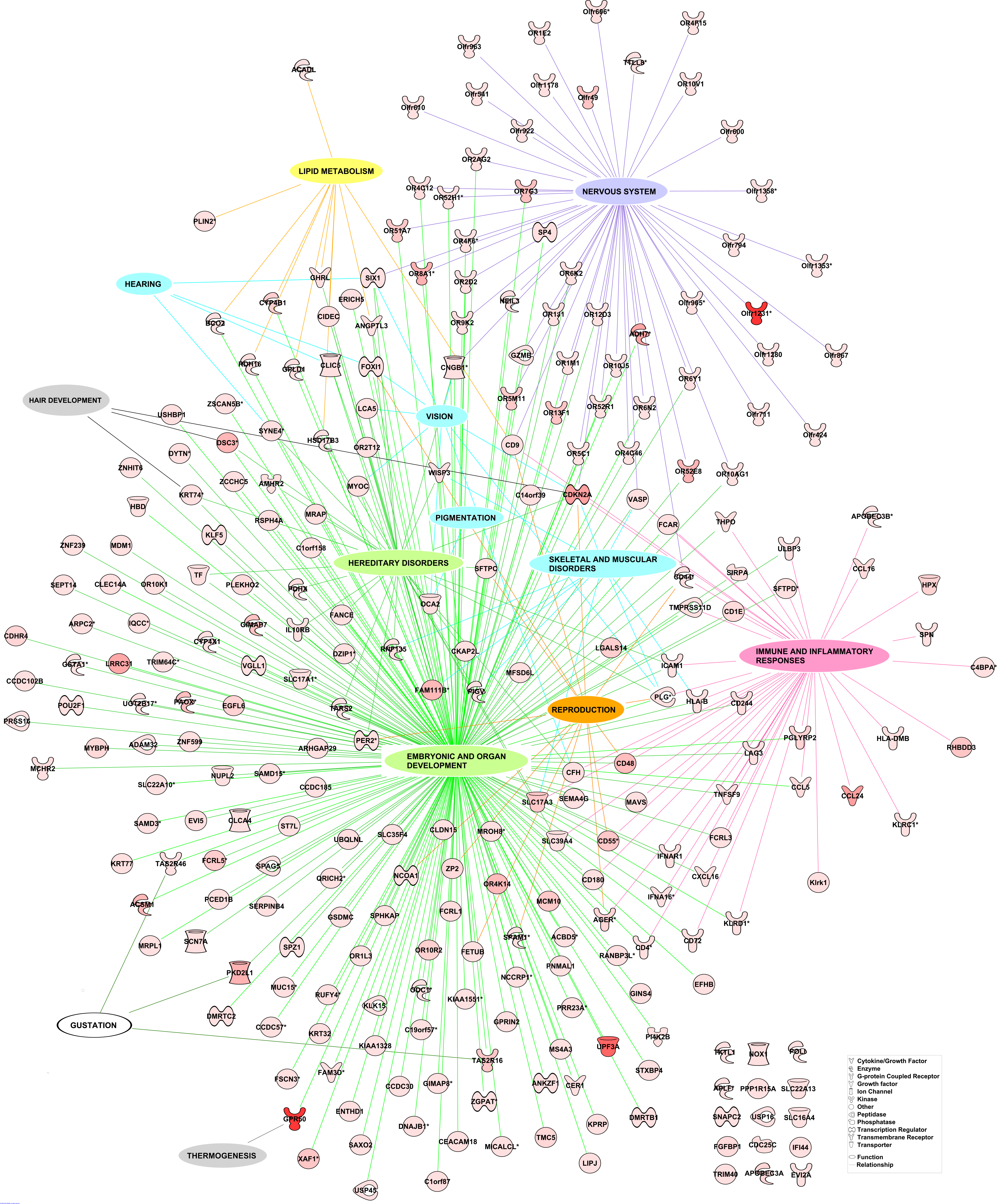
Representation of the genes with a Ka/Ks >1 connected to their key functions. Each global function and its links to the corresponding key genes are differently colored i.e. in purple for nervous system, blue for by-products of domestication, green for functions related to embryonic, organ development, organismal abnormality and hereditary disorders, pink for immune and inflammatory responses, orange for reproduction, grey for hair development and thermogenesis, yellow for lipid metabolism and white for gustation. Global functions contain genes belonging to several Ingenuity Pathway Analysis functional categories. Gene symbols are colored in red and color intensity is correlated to Ka/Ks value.

**Table 2:**
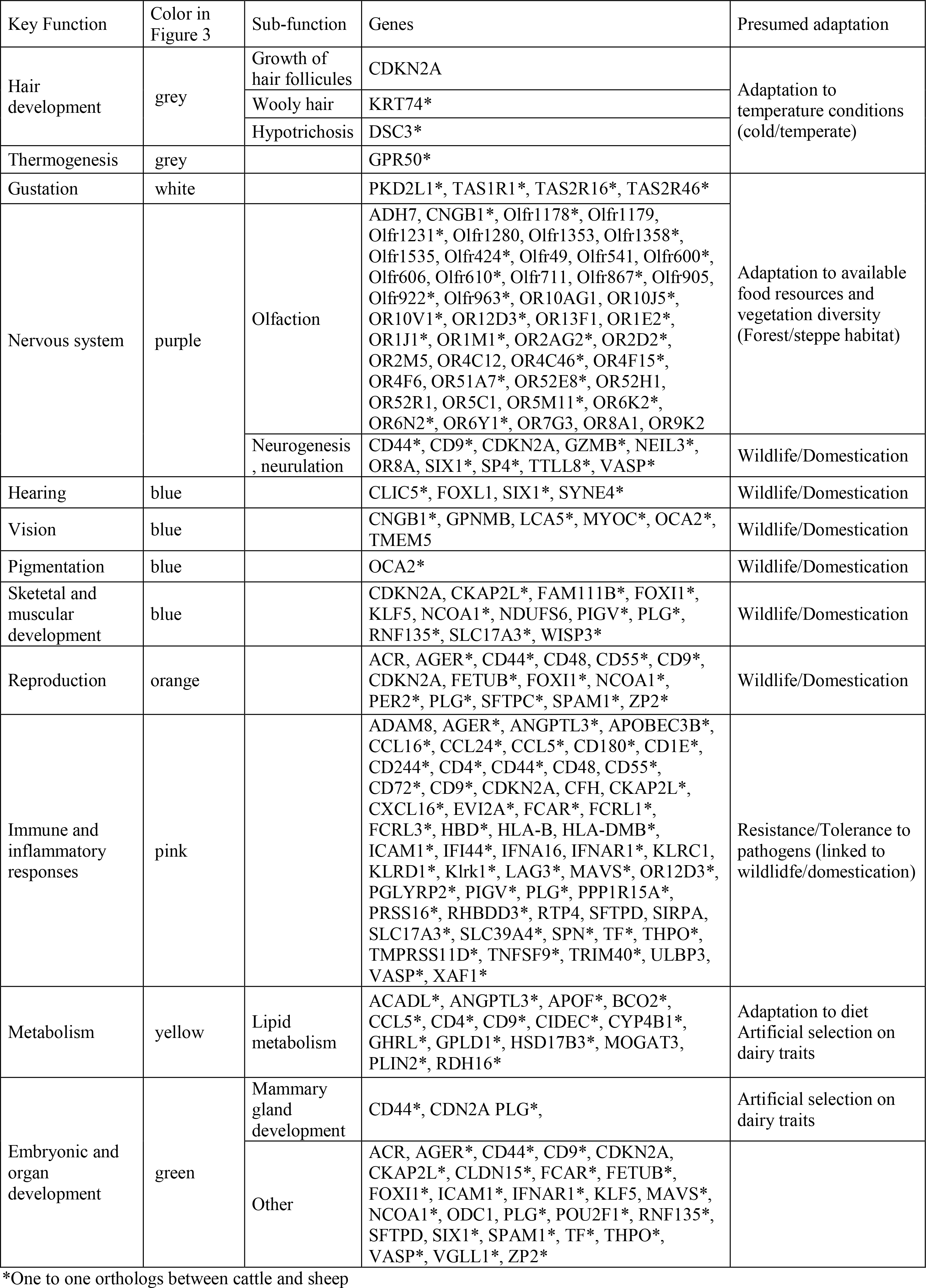
Main genes under selection listed by key functions and by presumed adaptation.

Among the genes under selection, some underlie obvious distinctive features between wisent and cattle providing in turn insights into the wisent adaptive history (Table 2, Figure 3 and Table S6). First, KRT74 and DCS3 that are involved in wooly hair development and hypotrichosis, respectively (Ayub, et al. 2009; Shimomura, et al. 2010) and GPR50 that plays a role in thermogenesis (Bechtold, et al. 2012) might be directly related to wisent adaptation to colder climatic conditions (Table 2, Table S4). Second, genes encoding olfactory and taste receptors are probably footprints of feeding behavior modifications, resulting *e.g.*, from food resource differences in forest *versus* steppe habitat (trees-shrubs *versus* grass). Third, the many genes related to immune and inflammatory responses (through their key role in activation, migration, binding, expansion and modulation of a broad range of immune cells, Table S5) may sign adaptation of wisent and bovine to different pathogen exposures. Fourth, several genes were found involved in lipid metabolism or mammary gland development (Table S5), likely the result of selection in cattle for improved milk production performances. More strikingly, the functional analysis of the genes under selection highlighted physiological functions associated to the domestication process (Figure 3, Table 2, Table S6), as expected from the close relatedness of wisent and domestic cattle. These functions underlie both key processes of nervous system (*e.g.*, neurogenesis, neurulation, remodeling of dendrites, differentiation of some neural cells) and by-product phenotypes of domestication (*e.g.*, skeletal and muscular disorders, hair and skin properties, vision, hearing, reproduction). Our analysis thus gives an empirical support to the unified explanation of the “domestication syndrome” in mammals (*i.e.*, the general combination of observed traits in domestic mammals) formulated by Wilkins and collaborators (Wilkins, et al. 2014) following the pioneering work by D.K. Belyaev and L. Trut (Belyaev 1979; Trut, et al. 2009). This explanation accords a central role to the neural crest through its developmental reduction as a result of the primary domestication pressure (*i.e.*, the taming of animals). Furthermore, because neural crest cells are also cellular precursors of many different cells (*e.g.*, osteocytes, chondrocytes, odontocytes and melanocytes), this reduction indirectly produces various secondary phenotypic changes (*e.g.*, skeletal and craniofacial morphological modifications or coat-colour changes). For instance, such an indirect relationship is well supported by studies on coat coloration in wild and domesticated animals (Cieslak, et al. 2011) and the description coat-colour associated mutations with pleiotropic effects (Reissmann and Ludwig 2013). Also, the numerous genes detected under selection and involved in immune response (Table 2 and Figure 3) might be indirectly related to the domestication process. During this process and afterwards, herding conditions may have incidentally lead to an increased pathogen exposure of domestic cattle due to proximity with individuals from the same or different species (Freeman, et al. 2008).

## Conclusions

To characterize the wisent demographic and adaptive histories, we carried out whole genome sequencing of two males from the Bialowieza lowland line. Although still considered as a vulnerable species, our results show that the conservation plan and subsequent management practices have been efficient to recover a reasonable amount of genetic variability that now compares to that observed in commercial cattle breeds. We further confirmed at the nucleotide level, the close relatedness of the wisent and cattle species without any evidence for recent gene flow (i.e., during historical times) between these two species. We estimated that the divergence between the bovine and wisent lineages occurred in the early Pleistocene through a speciation process involving limited gene flow, lasting from 1.7×10^−6^ to 850,000 YBP. We also found evidence for more recent secondary contacts, posterior to the *Bos taurus* and *Bos indicus* divergence (*ca*. 150,000 YBP), between the wisent and (European) taurine cattle lineages. Interestingly, whole individual genome based demographic inference highlighted contrasting patterns in both species that might be reminiscent of different adaptive strategies (habitat fragmentation versus migration) to survive Pleistocene glaciations. Our results are indeed in agreement with a scenario in which wisent survive in refugee pockets during glaciations (leading to habitat fragmentation) while aurochs migrate southward (and possibly eastward) where climate remained more temperate (Sommer and Nadachowski 2006). It is tempting to speculate that these two alternative strategies might have contributed to the survival of both species to the large mammals Quaternary extinction.Our results also show that wisent and aurochs display a similar trend towards extinction from the Last Glacial Maximum, 20,000 years ago that is concomitant to human population growth (spread of farming and increased hunting pressure), to climate warming and to vegetation changes (e.g., replacement of open-steppe by forests). For the wisent, in particular, the Holocene period was characterized by a tendency to shift towards forested habitats imposing a significant diet change (Kerley, et al. 2012; Bocherens, et al. 2015). These new constraints left footprints at the genomic level as illustrated by the genes that we found under selection when comparing the wisent and bovine lineages, i.e., genes involved in feeding behavior and in adaptation to temperature conditions and to pathogen exposure. Conversely, wisent being the closest extant wild relative species of domestic cattle, several of the genes under selection could be related to the adaptive response to cattle domestication via their implication in nervous system development and in the expression of by-product domestication phenotypes. Strikingly, this result is in line with unified explanation of the domestication syndrome in mammals formulated by Wilkins *et al.* (2014) giving a shared developmental connection of these diverse traits via neural crest cells.

## Materials and Methods

### Sample origin

Genomic DNA of two male wisents, namely BBO_3569 and BBO_3574, was extracted from blood samples collected in 1991 in the Bialowieza forest. More precisely, BBO_3569 and BB0_3574 belong to the genetically isolated pure lowland line which originates from seven founders kept in zoo and private breeding centers and used for the species restoration program at the beginning of the 1920’s (Tokarska, et al. 2009). In 1952, 40 of their descendants that were born in captivity were re-introduced to the wild in the Polish part of the Białowieża forest, the so-called lowland line now including more than 2,000 individuals (i.e., about half of the world wisent population).

### High-throughput sequencing of two wisents

The Illumina TruSeq DNA sample preparation kit (FC-121-2001, Illumina Inc., San Diego,USA) was used according to the manufacturer’s protocol. Libraries were then validated on a DNA1000 chip on a Bioanalyzer (Agilent) to determine size and quantified by qPCR using the Kapa library quantification kit (KAPA) to determine concentration.

The cluster generation process was performed on cBot (Illumina Inc.) by using the Illumina Paired-End DNA sample preparation kit (FC-102-1001, Illumina Inc.). Both BBO individuals were further paired-end sequenced on the HiSeq 2000 (Illumina Inc.) using the SBS (Sequence By Synthesis) technique. Base calling was achieved by using the RTA software (Illumina Inc.). Reads of 100 bp from both sides of the fragments were thus obtained after this step. Quality control of the sequences was checked using the FastQC software.

### Mapping wisent sequencing reads onto the bovine genome

In total, 331,975,598 and 407,585,788 reads paired in sequencing were available for BBO_3569 and BBO_3574 respectively after Illumina Quality Check. After removal of sequencing adapters, these reads were mapped onto the UMD3.1 cattle (*Bos taurus*) reference genome assembly (Liu, et al. 2009) using default options of *aln* and *sampe* programs from the *bwa* (version 0.6.2) software package (Li and Durbin 2009). In total, 92.7% (respectively 92.2%) of the reads from the BBO_3569 (respectively BBO_3574) library were successfully mapped onto the bovine assembly, 94.5% (respectively 93.6%) of which being properly paired. Read alignments with a mapping quality Phred-score MAPQ<20 and PCR duplicates were further removed using the *view* (option *−q 20*) and *rmdup* programs from the *samtools* (version 0.1.19) software package (Li and Durbin 2009). The resulting bam files are available for download from the Sequence Read Archive repository (http://www.ncbi.nlm.nih.gov/sra) under the accession number SRP070526. It should be noted that mapping the wisent reads onto the preliminary Y-chromosome sequence from the BosTau7 assembly (available at http://www.genome.ucsc.edu/) lead to a high proportion of reads improperly paired and only about 12% of the BTAY sequence was covered with an unexpected high read coverage (>60X). We thus decided not to consider the chromosome Y in further analyses.

### Construction of individual wisent consensus genome sequences

For the analyses of BBO/BTA cattle divergence, consensus genome sequences were built for each individual by first generating a *mpileup* file considering each European bison *bam* alignment file separately and using the *mpileup* program from the *samtools* (version 0.1.19) software package (Li and Durbin 2009) run with −C 50 and −d 5000 options. For each BBO individual and at each position (in the bovine reference assembly), the retained consensus base was randomly sampled among the aligned bases after discarding those showing a Base Alignment Quality BAQ>30 (allowing to account for uncertainty resulting from small indels). Such a procedure was aimed at limiting biases towards the bovine reference base at BBO heterozygous sites. Finally, positions with a read depth DP<3 and DP>30 (i.e., three times the average individual read coverage), or for which the retained base was supported by less than 2 reads (to limit sequencing error biases) were treated as N (not called) in the consensus sequences.

### Whole genome sequence of four individual bovine genomes

Four bovine individual whole genome sequences were obtained from the 1000 bull genome projects data (Daetwyler, et al. 2014) stored in the NCBI *sra* archive website (http://www.ncbi.nlm.nih.gov/sra). More precisely, the four males selected were AAN_0037, HOL_0101, JER_0009 and SIM_0043 and belonged to the Angus, the Holstein, the Jersey and the Simmental European taurine breeds respectively and were sequenced at a roughly similar coverage (8.2X, 9.6X, 11X and 10X respectively) than the European bisons. As for the European bison sequencing data, reads with a MAQ<20 and PCR-duplicates were filtered out from the downloaded *bam* files using the *view* (option *−q 20*) and the *rmdup* programs from the *samtools* (version 0.1.19) software package (Li and Durbin 2009).

### Testing for recent gene flow events between the BBO and BTA lineages using the ABBA-BABA statistics

To count the number of sites with ABBA and BABA patterns across the BTA, BIN, BBO and OAR lineages, we first identified sites displaying a “BA” pattern by comparing the consensus sequences dervived from each individual BBO to the OAR reference genome assembly. To that end and following the same procedure and program options as the ones described above for the mapping of reads onto the UMD3.1 bovine genome assembly (see Results section and Table S1), we mapped the sequencing reads originating from the BBO_3569 and BBO_3574 European bisons onto the *OviAr3* OAR genome assembly (Jiang, et al. 2014). Based on the resulting *mpileup* alignment files, we then defined for each BBO individual a consensus base as described above (section “Construction of individual consensus genome sequences”) except that the upper read depth threshold was set to 20 (to account for the lower coverage when mapping reads onto the ovine genome as summarized in see Table S1). The “BA” sites were those at which the consensus base differed from the OAR reference. To further identify whether the “A” or “B” allele was present in the BTA and BIN genomes, the OAR sequences surrounding each of these sites were extracted (60 nt upstream and 60 nt downstream) and aligned onto the BTA UMD3.1 (Liu, et al. 2009) and BIN genome (Canavez, et al. 2012) assemblies using the *blat* (version v35×1) software with default options except for the minimum sequence identity that was set to 95% (Kent 2002). For a given comparison, sequences aligning to more than 10 positions were discarded from further analyses (otherwise, the alignment displaying the highest score was retained). Similarly, sites for which a base different from the “A” and “B” alleles was present in the BIN or BTA sequence or the 2 upstream and downstream flanking nucleotides did not perfectly match were although discarded. In total, 10,035,538 (respectively 11,533,956) “BA” sites could be called in both the BIN and BTA assemblies based on the BBO_3564 (respectively BBO_3574) genomic data. From the observed numbers of “ABBA” and “BABA”, we then computed the *D-statistics* and both their associated standard error and their corresponding Z-score (to assess whether *D* significantly differs from zero) using a Weighted Block Jacknife with 5 Mb blocks (Green et al., 2010; Durand et al., 2011). To further estimate the proportion of BBO ancestry in the BTA genome, we relied on the estimator proposed by Durand et al. (2011) here defined as 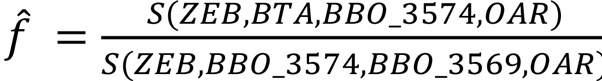, where S(P1,P2,P3,P4) represents the difference of the “ABBA” and “BABA” sites numbers assuming a ((P1;P2);P3);P4) phylogeny and estimated following a procedure similar to the one described above. As for the *D-statistic* estimates, the standard deviation of 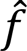 was computed using a Weighted Block Jacknife with 5 Mb blocks.

### Estimation of genetic heterozygosity from individual genomes

Genetic heterozygosities were estimated from the individual genome alignments onto the UMD 3.1 cattle assembly (i.e., based on the *mpileup* files described above) using *mlrho* version 2.8 (Haubold, et al. 2010). Note that *mlrho* actually implements a maximum likelihood estimator of the population mutation rate (*θ* = 4*N_e_μ_g_*), which fairly approximates heterozygosity under an infinite sites model (and providing *6* is small), while simultaneously estimating sequencing error rates and accounting for binomial sampling of parental alleles (Lynch 2008). For a given individual genome alignment, only sites covered by 3 to 30 reads (after discarding bases with a BAQ<25) were retained in the computation (Table S2).

### Population size history inference using the psmc

Effective population sizes in units of **4*N_e_μ_g_*** were estimated backward in time (in units of 2*μ_y_T*) based on individual whole genome sequences under the PSMC model using the program *psmc* (version 0.6.4) program (Li and Durbin 2011). Briefly, the analyzed individual whole genome sequences were obtained from the above described *mpileup* files (aligned onto the UMD3.1 cattle assembly) that were processed (individually) with *bcftools* (version 0.1.19) (Li and Durbin 2009) using the *−c* option. The resulting *vcf* files were further converted into *fastq* files using the *vcf2fq* utility of the *vcfutils.pl* program available in the *samtools* (version 0.1.19) suite (Li and Durbin 2009) discarding sites located less than 5 nucleotides from an indel (*−l5* option) and covered by less than 3 or more than 30 reads (*−d 3 −D 30* options). As originally described, the *psmc* program was finally run with default options except for the pattern of atomic time intervals *(-p* option) that were set to “4+25*2+4+6” on a reduced version of the (autosomal) genomes (*psmcfa* files), with consecutive sites grouped into consecutive bins of 100 nucleotides marked as “K” (at least one heterozygote), “T” (no heterozygote sites) or “N” (less than 90 called sites).

### Characterization of the divergence between BBO and BTA lineages under the coalHMM framework

For each BBO individual consensus genome sequence, genome alignment onto the UMD 3.1 bovine reference assembly (see the section “Construction of individual consensus genome sequences” above) was divided into 10 Mbp non overlapping segments. Segments mapping to the (bovine) X-chromosome or with more than 500,000 (5 %) missing information (i.e., an uncalled base in either the BTA or BBO sequence) were further discarded leading to a total of 123 and 177 available segments for the BBO_3569 and BBO_3574 alignments respectively. Maximum likelihood inference was then carried out independently for each segment using the scripts *isolation-model.py* (version 1.1) for the I-model, and the *initial-migration-model.py* (version 1.2) for the IM-model, from the *IMCoalHMM* software package (https://github.com/mailund/IMCoalHMM). More precisely, the parameters underlying the I-model include the split time *τ_D_* (in units of *μ_y_T*) and also the ancestral effective population size (in scaled units of ***θ* = 4*N_e_μ_g_***) and the recombination rate ρ (in scaled units of *μ_g_*) that both parameterize the underlying coalescent process. The parameters underlying the IM-model include the times *τ_1_* (completion of the split) and the IM period length *τ_M_* that both are units of *μ_y_T*, the migration rate *M* (in number of migration per substitution) and *θ* and ρ (as for the I-model). Note that the product *M* × (*θ*/2) corresponds to the estimated number of migrant per generation (in the usual units of **2*N_e_m*** where *m* is the net migration rate) over the IM period. Finally, for model comparison purposes, the Akaike Information Criterion (AIC) was computed as AIC_I_=2×(3−*l*) for the I-model and AIC_IM_=2×(5−*l*) for the IM-model where *l* represents the estimated log-likelihood of the corresponding model.

### Detection of genes under selection

Based on the alignment of the BBO_3569 and BBO_3574 individual sequences onto the bovine UMD 3.1 reference genome assembly (*mpileup* file described above), a consensus BBO sequence was derived for all the n=22,091 (bovine) ENSEMBL protein-coding sequences as defined in the UCSC genome database (http://genome.ucsc.edu/). At a given position, the retained consensus base corresponded to the most represented one among all the aligned reads from both the BBO_3569 and BBO_3574 individuals and after discarding bases with a BAQ<30. Positions with a read depth DP<3 and DP>60 (three times the average coverage of BTA genome assembly by the combined BBO_3569 and BBO_3574 read sequencing data) were treated as N (not called) in the consensus sequences. BBO consensus protein sequences with more than 10% “N” were discarded from further analysis leading to a total of n=19,372 remaining BBO/BTA alignments (with 1.11% “N” on average). For 1,306 (6.74%) of these, no nucleotide differences could be observed between the obtained BBO consensus sequence and the BTA one. Nonsynonymous (Ka) and synonymous (Ks) substitution rates were then computed using *KaKs_calculator v1.2* (Zhang, et al. 2006) assuming a MLWL model to estimate the number of synonymous and nonsynonymous sites (Zhang, et al. 2006) allowing the estimation of the *Ka/Ks* ratio for a total of 17,073 protein coding sequences.

### Functional annotation of candidate genes under positive selection and gene network analysis

Among the 17,073 Ensembl transcripts ID corresponding to the protein coding sequences for which a *KaKs* ratio was estimated, 873 transcripts with a *KaKs* ratio above one were considered under positive selection.

Functional annotation of these transcripts and networks analyses were carried out with the Ingenuity Pathway Analysis software (IPA, Ingenuity®Systems, www.ingenuity.com). Among the 873 transcripts with a *KaKs* ratio above one, 481 transcript ID were mapped to the Ingenuity Pathway Knowledge Base (IPKB) and were representative of 425 individual genes ready for analysis (Table S3). A total of 405 of these transcripts were further identified as one to one orthologs between cattle and sheep, using Ensembl orthology information (http://www.ensembl.org/biomart/). The top significant functions and diseases (p-value<0.05) were obtained by comparing functions associated with the 425 (among which 359 were one to one orthologs between cattle and sheep) genes against functions associated with all genes in our reference set (14,465 transcripts mapped in IPKB from the 17,073 transcripts ID that were analyzed) using the right-tailed Fisher's exact test. A total of 262 genes (among which 151 were one to one orthologs between cattle and sheep) participated to the significant functions thus determined (Table S5).

Among the 425 genes, 367 genes (among which 324 were one to one orthologs between cattle and sheep) were included in network analyses. For each network that contains at most 140 molecules, a score S was computed based on a right-tailed Fisher exact test for the overrepresentation of the genes with a *KaKs* ratio>1 (S=−log(p-value)). A network was considered as significant when S>3.

## Acknowledgements and funding information

The authors wish to thank Anna Madeyska-Lewandowska (Private Veterinary School, Warsaw, Poland) that provided the *Bison bonasus* samples in the early 1990's as part of a collaborative research project on the characterization of casein variants. We are also grateful to the four anonymous reviewers for their very helpful and constructive comments. This work was supported by an INRA (Institut National de La Recherche Agronomique)-Animal Genetics Department Grant (AFROSEQ project).

Sequencing data (bam files) are available for download from the Sequence Read Archive repository (http://www.ncbi.nlm.nih.gov/sra) under the accession number SRP070526.

## SUPPLEMENTARY TABLE LEGENDS

**Table S1: Mapping statistics of the European bison high throughput sequencing reads onto the UMD3.1 bovine reference genome and *OviAr3* ovine reference genome assemblies.**

**Table S2: Maximum likelihood estimates of the (autosomal) heterozygosities (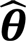) and sequencing error rates (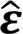) for the BBO_3569 and BBO_3574 individual European bison genomes and the AAN_0037, JER_0009, HOL_0101 and SIM_0043 individual genomes belonging to the Angus, Jersey, Holstein and Simmental European cattle breeds respectively.** The table also gives the number of sites available in the estimation procedure (i.e., covered by 10 to 100 read nucleotides after discarding bases with a BAQ<25)

**Table S3: List of the 481 Ensembl transcript ID with a Ka/Ks >1 (from 1.003 to 8.289) mapped in the Ingenuity Pathway Knowledge Base and of their corresponding gene symbol and Entrez gene name.** One to one orthologs between cattle and sheep are indicated.

**Table S4: List of the top five significant diseases and biological functions in each main functional categories obtained using Ingenuity Pathway Analysis for the 481 (405) transcripts with a Ka/Ks >1 (one to one orthologs between bovine and sheep).**

**Table S5: List of the significant functional categories with their corresponding functional annotation, p-value and annotated molecules.**

**Table S6: List of the significant networks including genes with a Ka/Ks >1, obtained with the Ingenuity Pathway Analysis software.**

### SUPPLEMENTARY FIGURE LEGENDS

**Figure S1: Regional nucleotide divergence between the cattle genome and the BBO_3569 (A) and BBO_3574 (B) European bison genome sequences over contiguous 10-Mb segments covering the 29 bovine autosomes and the X-chromosome.**

**Figure S2: Distribution of the heterozygosity estimates across the 29 bovine autosomes based on the BBO_3569 and BBO_3574 individual European bison genomes and for the AAN_0037, JER_0009, HOL_0101 and SIM_0043 individual genomes belonging to the Angus, Jersey, Holstein and Simmental European cattle breeds respectively.**

**Figure S3: Backward in time (in units of 2*μ_y_T*) estimates of the (scaled) effective population sizes (in units of 4*N_e_μ_g_*) from the analyses of the BBO_3569 and BBO_3574 individual wisent genomes and the AAN_0037, HOL_0101, JER_0009 and SIM_0043 bull bovine genomes**

**Figure S4: Boostrap confidence intervals of the population size histories inferred from the two wisent and the four bovine genomes under the PSMC model. For each individual genome, the dark red curve represents the direct estimation (as in Figure 1A) while the 100 orange curves represent estimated history obtained on bootstrap genome samples (Li and Durbin 2011).**

**Figure S5: Comparison of the population histories inferred by the PSMC algorithm from the two wisent genomes derived by either aligning the underlying sequencing reads onto the UMD 3.1 cattle genome assembly (in black) or the consensus BBO reference assembly (in red).**

**Figure S6: Distribution of the Ka/Ks estimates obtained for all the BBO/BTA protein sequence alignments.**

**Figure S7. List of genes with a Ka/Ks ratio>1 participating to the six overlapping significant networks.** Gene symbols are colored in red and color intensity is correlated to Ka/Ks value.

